# Divergent *Plasmodium* kinases drive MTOC, kinetochore and axoneme organisation in male gametogenesis

**DOI:** 10.1101/2024.09.22.613647

**Authors:** Ryuji Yanase, Mohammad Zeeshan, David J. P. Ferguson, Robert Marcus, Declan Brady, Andrew R. Bottrill, Anthony A. Holder, David S. Guttery, Rita Tewari

## Abstract

Sexual development and male gamete formation of the malaria parasite in the mosquito midgut is governed by rapid endomitosis in the activated male gametocyte. This process is highly regulated by protein phosphorylation, specifically by three divergent male-specific protein kinases (PKs): CDPK4, SRPK1 and MAP2. Here, we localise each PK during rapid male gamete formation using live-cell imaging, identify their putative substrates by immunoprecipitation, and determine the morphological consequences of their absence using ultrastructure expansion and transmission electron microscopy. Each PK has a distinct location in either the nuclear or cytoplasmic compartment. Protein interaction studies revealed that CDPK4 and MAP2 interact with key drivers of rapid DNA replication, while SRPK1 is involved in RNA translation. The absence of each PK results in severe defects in either microtubule organising centre (MTOC) organisation, kinetochore segregation or axoneme formation. This study reveals the crucial role of these PKs during endomitosis in formation of the flagellated male gamete and uncovers some of their potential substrates that drive this process.

**Summary blurb:** This study reveals how Plasmodium kinases regulate MTOC, axoneme and kinetochore organisation in male gametogenesis, providing key insights into potential targets for malaria transmission control.

## Introduction

Mitotic cell division of the malaria parasite, Plasmodium, is characterised by two forms of atypical, closed mitosis. The first is schizogony in hepatocytes and red blood cells of the mammalian host and sporogony within oocysts in the mosquito gut. The second is in development of activated male gametocytes within the mosquito midgut during sexual development to form eight haploid, flagellated, motile male gametes. Closed mitosis during male gametogenesis involves three rapid rounds of DNA replication over 8 to 10 min in an endomitotic division to produce an octoploid nucleus. Only once all rounds of nuclear division are complete does exflagellation occur to produce male gametes (Sinden, 2015; Guttery et al, 2022).

Reversible protein phosphorylation, catalysed by protein kinases (PKs) and protein phosphatases (PPs), is a ubiquitous, highly-conserved mechanism of protein activation/deactivation that is known to regulate mitosis in Eukaryota (Nilsson, 2019; Strumillo et al, 2019; Fulcher & Sapkota, 2020). Protein components of key structures in mitosis, such as microtubule-organising centres (MTOCs), spindles, and kinetochores, are subject to spatiotemporal regulation mediated by the coordinated activity of various PKs and PPs to enable accurate cell cycle control and chromosome segregation (Ito & Bettencourt-Dias, 2018; Ong et al, 2020; Vagnarelli, 2021; Kucharski et al, 2022). In the rodent malaria parasite Plasmodium berghei (Pb), several studies have highlighted three male-specific PKs, which are defined as essential during male gametogenesis but not at the asexual blood stage. These are calcium-dependent protein kinase-4 (CDPK4), a serine/arginine-rich protein kinase (SRPK1) and mitogen-activated protein kinase-2 (MAP2; Billker et al., 2004; Guttery et al., 2024; Tewari et al., 2005; Tewari et al., 2010). The deletion of each ablates exflagellation (male gamete release) and completely blocks sexual reproduction and parasite transmission through the mosquito. However, the deletion of each PK has little to no effect on the development of the parasites during the asexual blood stage (Billker et al, 2004; Tewari et al, 2005, 2010). Each PK has been postulated to act at a distinct point: CDPK4 is considered to be a master regulator, initiating assembly of the pre-replicative complex and formation of the first mitotic spindle within the first 18 seconds following gametocyte activation (Billker et al, 2004; Fang et al, 2017; Invergo et al, 2017); SRPK1 is required for subsequent DNA replication (Invergo et al, 2017), and MAP2 is required for cytokinesis and axoneme motility (Tewari et al, 2005; Guttery et al, 2012).

Throughout DNA replication the MTOC is crucial to its completion (Ito & Bettencourt-Dias, 2018; Guttery et al, 2022). The MTOC has different names in different species; for example, in yeast it is known as the spindle pole body (SPB), in humans the centrosome, and in ciliated cells the basal body (Seybold & Shiebel, 2013; Vaughan & Gull, 2016; Vertii et al, 2016; Ito & Bettencourt-Dias, 2018). As an anchor for mitotic spindles, the MTOC is assembled, duplicated and separated at each round of mitosis (Wu & Akhmanova, 2017), which during Plasmodium male gametogenesis results in the formation of eight flagellated male gametes (Guttery et al, 2022). Several PKs are associated with MTOC function (Vertii et al, 2016), for example in humans and Drosophila, Aurora kinase A and polo-like kinase 1 (Plk1) are recruited to the spindle poles and play a crucial role in spindle assembly and mitotic progression (Barr & Gergely, 2007). Also, never in mitosis (NIMA)-like kinases (NEKs) and cyclin-dependent kinases (CDKs) are known regulators of MTOC biology in diverse organisms, including Aspergillus, plants, yeast and human cells (Joubes et al, 2000; Elserafy et al, 2014; Fry et al, 2017; Panchal & Evan Prince, 2023). In apicomplexan parasite Toxoplasma gondii (T. gondii), Aurora, NEK, and CDK-related kinase (CRK) have been shown to localise to mitotic structures and play crucial roles in cell cycle regulation (Chen & Gubbels, 2013; Suvorova et al, 2015; Gaji et al, 2021). Plasmodium NEK1 kinase was shown recently to be an important component of MTOC organisation and an essential regulator of chromosome segregation during male gamete formation (Zeeshan et al, 2024). Plasmodium CRK5 that is another male-specific divergent PK, was also shown recently to be a critical kinase, regulating male gametogenesis (Balestra et al, 2020; Kumar et al, 2022). Aurora kinase B is a known essential regulator of chromosome segregation and cytokinesis in organisms ranging from yeast to humans (Honda et al, 2003; Carmena et al, 2009, 2015; Hadders & Lens, 2022), and the highly divergent aurora kinase paralogue Plasmodium ARK2 has been shown to be essential for spindle dynamics and chromosome segregation during male gametogenesis (Zeeshan et al, 2023). However, the involvement of other divergent male-specific kinases in chromosome segregation in Plasmodium is not well understood.

Both CDPK4 and SRPK1 may play early roles in the homeostasis of the gametocyte bipartite MTOC and axoneme formation (Rashpa & Brochet, 2022; Zeeshan et al, 2022a). In CDPK4 deletion mutants, axoneme formation is severely impaired, with only short, incomplete axonemes being formed, while SRPK1 mutant parasites have incorrectly positioned basal bodies (Rashpa & Brochet, 2022). In contrast, MAP2 mutant lines form normal axonemes but have a severe defect in chromosome condensation (Guttery et al, 2012). It remains unclear which axoneme-associated proteins are phosphorylated by these PKs, and what their specific roles are during axoneme formation.

We aimed to localise each PK in relation to MTOC assembly, axoneme formation, and chromosome segregation, as well as identify some of their interacting partner proteins – particularly for MAP2 and SRPK1 for which little is known. Using a combination of live-cell imaging, protein identification by proteomics and ultrastructure analysis by expansion and electron microscopy, we show that PbCDPK4 and PbSRPK1 are diffusely distributed throughout the parasite cell, with a slightly stronger punctate cytoplasmic localisation during schizogony and male gametogenesis, whereas PbMAP2 is only in the nucleus and during male gametogenesis. The three PKs interact with each other and with key regulators of MTOC biology and male gametogenesis. The ultrastructure analysis revealed that their absence affects MTOC formation and separation, kinetochore dynamics, and axoneme development.

## Results and Discussion

### CDPK4, SRPK1 and MAP2 have different spatiotemporal locations during schizogony and male gametogenesis

Despite several studies highlighting the essential roles of CDPK4, SRPK1 and MAP2 in male gametogenesis in P. berghei (Billker et al, 2004; Tewari et al, 2010), little is known about their location during the endomitotic stages of cell division (i.e., schizogony, sporogony and male gametogenesis). To examine the spatiotemporal expression of CDPK4, SRPK1 and MAP2, we generated transgenic P. berghei parasite lines expressing the genes modified to code for a C-terminal GFP tag. An in-frame gfp coding sequence was inserted at the 3′ end of the endogenous gene locus (CDPK4 – PBANKA_0615200; SRPK1 – PBANKA_0401100; MAP2 – PBANKA_0933700), using single crossover homologous recombination (Figure S1A, Table S1 for primers), and successful insertion was confirmed by diagnostic PCR (Figure S1B, Table S1). Western blot analysis of schizont (for CDPK4 and SRPK1) and gametocyte (for MAP2 as MAP2 seems to be barely expressed in asexual blood stages) protein extracts, using an anti-GFP antibody, revealed major bands at 90 kDa for CDPK4-GFP, 183 kDa for SRPK1-GFP, 90 kDa for MAP2-GFP, and 29 kDa for unfused GFP (WT-GFP); the expected sizes for each. Each GFP-tagged PK was also observed by fluorescence in activated gametocytes (Figure S1C).

Live-cell fluorescence imaging of P. berghei asexual blood stages revealed a diffuse cytoplasmic distribution of CDPK4-GFP in both the schizonts and merozoites. In contrast SRPK1-GFP showed weak diffuse cytoplasmic expression with a strong focus at the location of the nuclear pole and towards the apex of the forming or mature merozoites (Figure 1A). In contrast, no MAP2-GFP fluorescence was observed in asexual development. During male gametogenesis, CDPK4-GFP had a diffuse location in both cytoplasm and nucleus, along with a distinct focus in the cytoplasm six min post-activation (Figure 1B, arrows). At the early stage of male gametogenesis (30 s to 6 min post-activation), up to two foci of CDPK4-GFP were observed, whereas at the late stage (>6 min post-activation, more than three foci were sometimes observed. SRPK1-GFP was located in the cytoplasm, concentrically surrounding the nucleus, confirming previous observations in P. falciparum (Pf; Dixit et al, 2010; Kumar et al, 2021). MAP2-GFP had a nuclear location in activated male gametocytes (Figure 1B).

**Figure 1.**
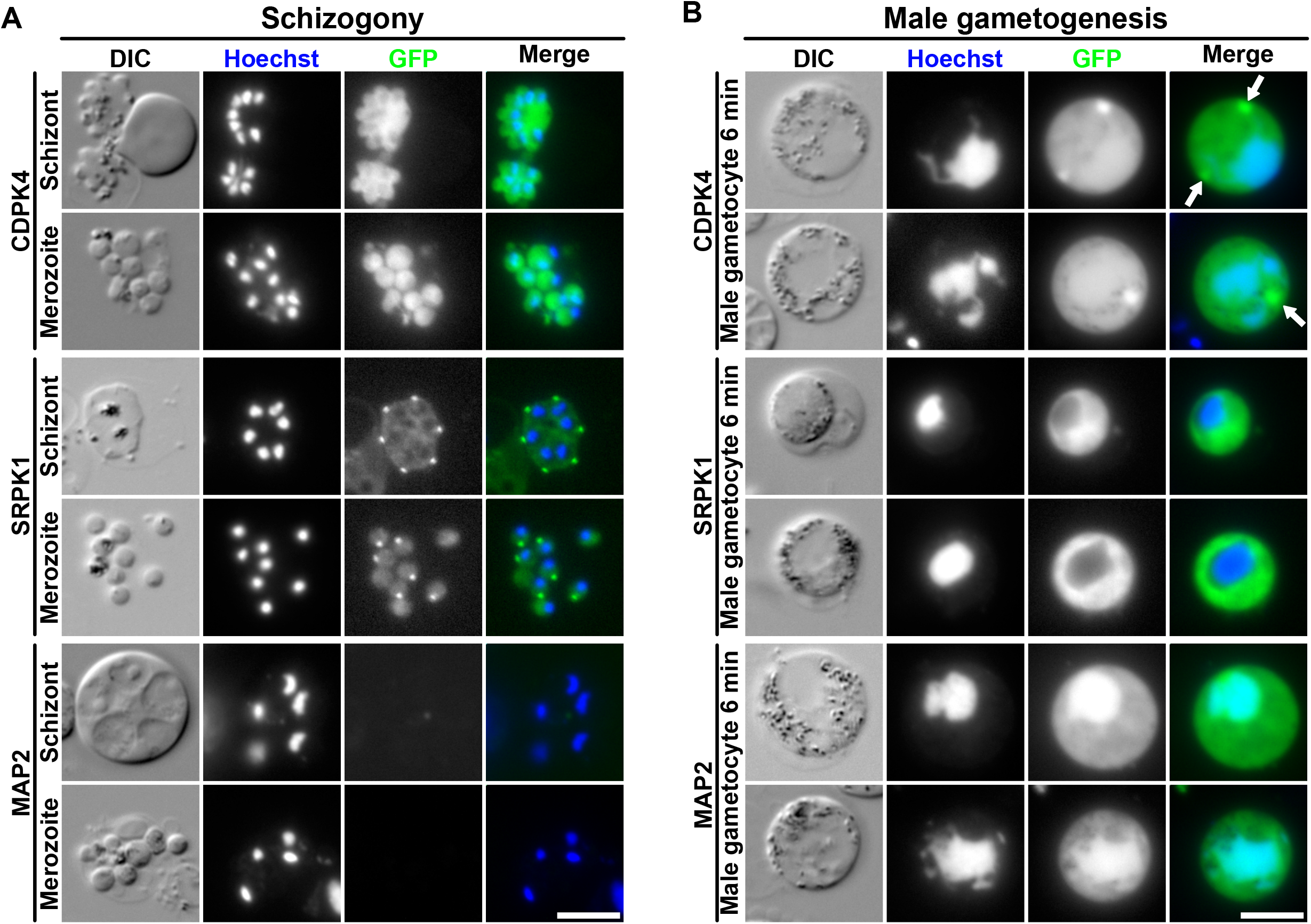
Fluorescence localisation in PK-tagged lines during asexual and sexual stages. (A) Localisation of each GFP-tagged PK in schizonts or merozoites. DIC: differential interference contrast. Merge shows Hoechst (blue) and GFP (green) signal. Scale bar = 5 µm. (B) Localisation of each GFP-tagged PK in activated gametocytes 6 min post-activation. Merge shows Hoechst (blue) and GFP (green) signals. White arrows indicate CDPK4-GFP concentrated focus in the cytoplasm. Scale bar = 5 µm.

Because these PKs are essential for axoneme development and are located at distinct regions of the parasite cell (in particular the distinct CDPK4-GFP focus in male gametocytes), we examined their co-localisation with markers of MTOC biology, including chromosome segregation (kinetochore protein, NDC80; Zeeshan et al., 2020) and basal body formation (cytoplasmic axonemal protein, kinesin-8B; Zeeshan et al., 2019), which were C-terminally tagged with mCherry (mCh). No co-localisation was observed between CDPK4-GFP, SRPK1-GFP or MAP2-GFP and NDC80-mCh during male gametogenesis (Figure 2A), or between SRPK1-GFP and NDC80-mCh during schizogony (Figure S2), suggesting that these PKs do not directly interact with kinetochores during chromosome condensation. However, kinesin-8B-mCh appeared as a ring around SRPK1-GFP, suggesting that SRPK1 may be interacting with axoneme proteins or proteins involved in their assembly (Figure 2B). CDPK4-GFP did not co-localise with kinesin-8B-mCh at any point (Figure 2B). This was surprising, since altered kinesin-8B phosphopeptide abundance has been observed in CDPK4 mutants (Fang et al, 2017; Kumar et al, 2021). A MAP2-GFP/kinesin-8B-mCh cross was not generated since these proteins are primarily located in different subcellular compartments.

**Figure 2.**
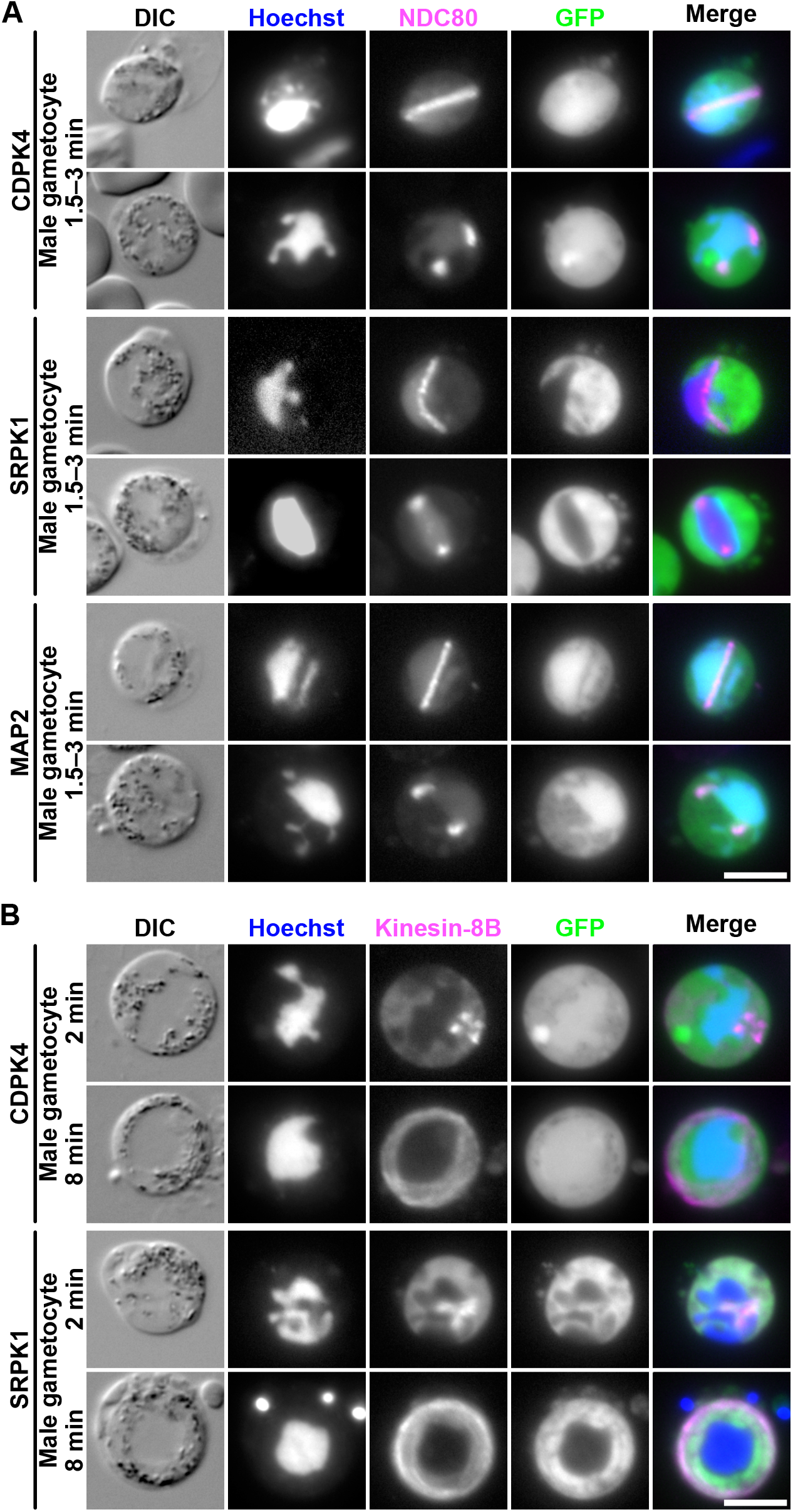
Co-localisation of PKs with kinetochore and basal body/axoneme markers. (A) Co-localisation of each GFP-tagged PK with mCherry (mCh)-tagged NDC80 in gametocytes, 1.5-3 min post-activation. DIC: differential interference contrast. Merge shows Hoechst (blue), NDC80-mCh (magenta) and GFP (green). Scale bar = 5 µm. (B) Co-localisation of each GFP-tagged PK with mCh-tagged kinesin-8B in gametocytes 2-8 min post-activation. Merge shows Hoechst (blue), kinesin-8B-mCh (magenta) and GFP (green). Scale bar = 5 µm.

### CDPK4, SRPK1 and MAP2 interact with each other and key regulators of male gametogenesis

Previous studies have highlighted differential protein phosphorylation, detected by altered abundance of phosphopeptides in protease digests, of cellular proteins associated with distinct cell-cycle events in deletion mutants of PbCDPK4 and PbSRPK1 (Fang et al, 2017; Invergo et al, 2017). In addition, SRPK1 was suggested to be phosphorylated in a CDPK4-dependent manner (Invergo et al, 2017). Fang et al., 2017 identified CDPK4-interacting proteins and CDPK4-dependent phosphorylation, using immunoprecipitation and mass spectrometry-based phosphoproteomics. However, the protein substrates of SRPK1 and MAP2 during male gametogenesis are largely unknown. We analysed the interactome of each PK in P. berghei gametocyte lysates to identify substrates that may regulate male gametogenesis.

In triplicate experiments, we immunoprecipitated with anti-GFP antibody CDPK4-GFP, SRPK1-GFP, MAP2-GFP and WT-GFP (as a control) from lysates of cells, 6 min post-activation of gametocytes. Immunoprecipitates were digested, and the resultant peptides analysed by mass spectrometry to identify component proteins. Potential interacting partners for each PK were identified as those with unique peptide sequences present in all three biological replicates, but completely absent from WT-GFP control samples.

A total of 108, 201 and 162 proteins were immunoprecipitated together with CDPK4-GFP, SRPK1-GFP and MAP2-GFP, respectively (Figure 3A, Table 1, Table S2-4). Gene ontology (GO) analysis of CDPK4- and MAP2-GFP partners highlighted several components of the replisome and DNA replication (Figure 3B, D, Table S5, S7). Of note, in the CDPK4-GFP line all five known Plasmodium origin recognition complex (ORC) subunits and CDC6 (components of the prereplicative complex; Chou et al., 2021) were present (Table 1, Table S2), along with the catalytic subunits of DNA polymerase alpha and epsilon. However, no mini-chromosome maintenance (MCM) proteins were detected, unlike in the MAP2-GFP samples, in which MCM3 to 7 were detected (Table 1, Table S4). These data are consistent with a role of CDPK4 in activation of the prereplicative complex (Invergo et al, 2017), and this role may explain the complete lack of DNA replication in CDPK4 deletion mutants (Tewari et al, 2010). In higher eukaryotes, phosphorylation of ORCs is facilitated by CDKs (Lee et al, 2012); whereas Cdc7-Dbf4 PK (DDK) promotes assembly of a stable Cdc45-MCM complex (Sheu & Stillman, 2006). No homologues of either Cdc7, Dbf4 or Cdc45 have been identified in the Plasmodium genome and therefore, we suggest that MAP2 performs the function of DDK. MCM2, 4 and 7 were also co-precipitated with SRPK1-GFP (Table 1, Table S3). Since phosphorylation of MCM2 is critical for its loading onto chromatin (Chuang et al, 2009), it is possible that overall CDPK4 is critical for association of the ORC with chromatin and the replication origin, whereas SRPK1 and MAP2 facilitate activation of the MCM complex. Altered phosphopeptides were only observed in SRPK1 deletion mutants for MCM5 and in CDPK4 mutants for ORC1 (Invergo et al, 2017), so this hypothesis differs somewhat from that of Fang et al., 2017, who identified CDPK4 as part of the MCM complex in non-activated gametocytes. However, in this study, gametocytes were analysed 6 min post-activation, a time point at which MCM function may be regulated by SRPK1 or MAP2, rather than CDPK4.

**Table 1.** Binding partners that are potential substrates of male-specific protein kinases during male gametogenesis List of proteins interacting with CDPK4-GFP, SRPK1-GFP and MAP2-GFP during male gametogenesis 6 min post-activation. Average unique peptides represent average across 3 biological replicates.

|  |  |  |  | Average no. Unique peptides |  |  |
| --- | --- | --- | --- | --- | --- | --- |
|  | *P. berghei* gene ID | Product description | Gene names | CDPK4-GFP | SRPK1-GFP | MAP2-GFP |
| Reversible protein phosphorylation | PBANKA_0615200 | Calcium-dependent kinase 4 | CDPK4 | 42 | 6 | 10 |
|  | PBANKA_0401100 | Serine/threonine protein kinase, putative | SRPK1 | 5 | 66 | 0 |
|  | PBANKA_0933700 | Mitogen activated protein kinase 2 | MAP2 | 5 | 0 | 28 |
|  | PBANKA_1227400 | Serine/threonine protein phosphatase 2B catalytic subunit A | CNA | 3 | 0 | 0 |
|  | PBANKA_1131900 | Serine/threonine protein phosphatase 5 | PP5 | 4 | 0 | 0 |
|  | PBANKA_0607400 | Protein phosphatase PPM11, putative | PPM11 | 20 | 0 | 13 |
| DNA replication | PBANKA_0602000 | Origin recognition complex subunit 1, putative | ORC1 | 24 | 0 | 0 |
|  | PBANKA_0803000 | Origin recognition complex subunit 2, putative | ORC2 | 23 | 0 | 8 |
|  | PBANKA_0513900 | ORC3 domain-containing protein, putative | ORC3 | 23 | 0 | 5 |
|  | PBANKA_1348800 | Origin recognition complex subunit 4, putative | ORC4 | 16 | 0 | 0 |
|  | PBANKA_0312500 | Origin recognition complex subunit 5, putative | ORC5 | 19 | 0 | 6 |
|  | PBANKA_1102900 | Cell division control protein 6, putative | CDC6 | 12 | 0 | 9 |
|  | PBANKA_1024900 | DNA replication licensing factor MCM2, putative | MCM2 | 0 | 5 | 0 |
|  | PBANKA_1241800 | DNA replication licensing factor MCM3, putative | MCM3 | 0 | 0 | 21 |
|  | PBANKA_1415600 | DNA replication licensing factor MCM4, putative | MCM4 | 0 | 3 | 27 |
|  | PBANKA_0610200 | DNA replication licensing factor MCM5, putative | MCM5 | 0 | 0 | 18 |
|  | PBANKA_1131600 | DNA replication licensing factor MCM6, putative | MCM6 | 0 | 0 | 17 |
|  | PBANKA_0803100 | DNA replication licensing factor MCM7, putative | MCM7 | 0 | 3 | 22 |
| Translation/mRNA splicing | PBANKA_1428700 | eukaryotic translation initiation factor 3 subunit A, putative | eIF3A | 0 | 14 | 0 |
|  | PBANKA_1232500 | eukaryotic translation initiation factor 3 subunit B, putative | eIF3B | 0 | 9 | 0 |
|  | PBANKA_0604800 | eukaryotic translation initiation factor 3 subunit C, putative | eIF3C | 0 | 9 | 0 |
|  | PBANKA_1206100 | eukaryotic translation initiation factor 3 subunit D, putative | eIF3D | 0 | 13 | 0 |
|  | PBANKA_1242800 | eukaryotic translation initiation factor 3 subunit E, putative | eIF3E | 4 | 3 | 3 |
|  | PBANKA_0715200 | eukaryotic translation initiation factor 3 subunit G, putative | eIF3G | 0 | 4 | 0 |
|  | PBANKA_0614500 | eukaryotic translation initiation factor 3 subunit I, putative | eIF3I | 0 | 6 | 0 |
|  | PBANKA_0720300 | eukaryotic translation initiation factor 3 subunit M, putative | eIF3M | 0 | 5 | 0 |
|  | PBANKA_0711900 | Heat shock protein 70 | HSP70 | 0 | 37 | 0 |
|  | PBANKA_0805700 | Heat shock protein 90, putative | HSP90 | 0 | 32 | 0 |
|  | PBANKA_0610900 | HSP40, subfamily A, putative | HSP40 | 0 | 27 | 0 |
| Chromosome organisation/MTOC | PBANKA_1416900 | Structural maintenance of chromosomes protein 2 | SMC2 | 7 | 0 | 11 |
|  | PBANKA_1108700 | Structural maintenance of chromosomes protein 4 | SMC4 | 12 | 0 | 18 |
|  | PBANKA_0917500 | Structural maintenance of chromosomes protein 1, putative | SMC1 | 0 | 7 | 8 |
|  | PBANKA_0716000 | Structural maintenance of chromosomes protein 3, putative | SMC3 | 0 | 6 | 5 |
|  | PBANKA_0917400 | Armadillo repeat protein PF16 | PF16 | 0 | 6 | 8 |
|  | PBANKA_0202700 | Kinesin-8B, putative | Kinesin-8B | 0 | 12 | 14 |
|  | PBANKA_1458300 | Kinesin-13, putative | Kinesin-13 | 6 | 0 | 6 |
|  | PBANKA_1458800 | Kinesin-15, putative | Kinesin-15 | 0 | 0 | 8 |
|  | PBANKA_1310400 | Centrin-2, putative | Centrin-2 | 0 | 6 | 0 |

**Figure 3.**
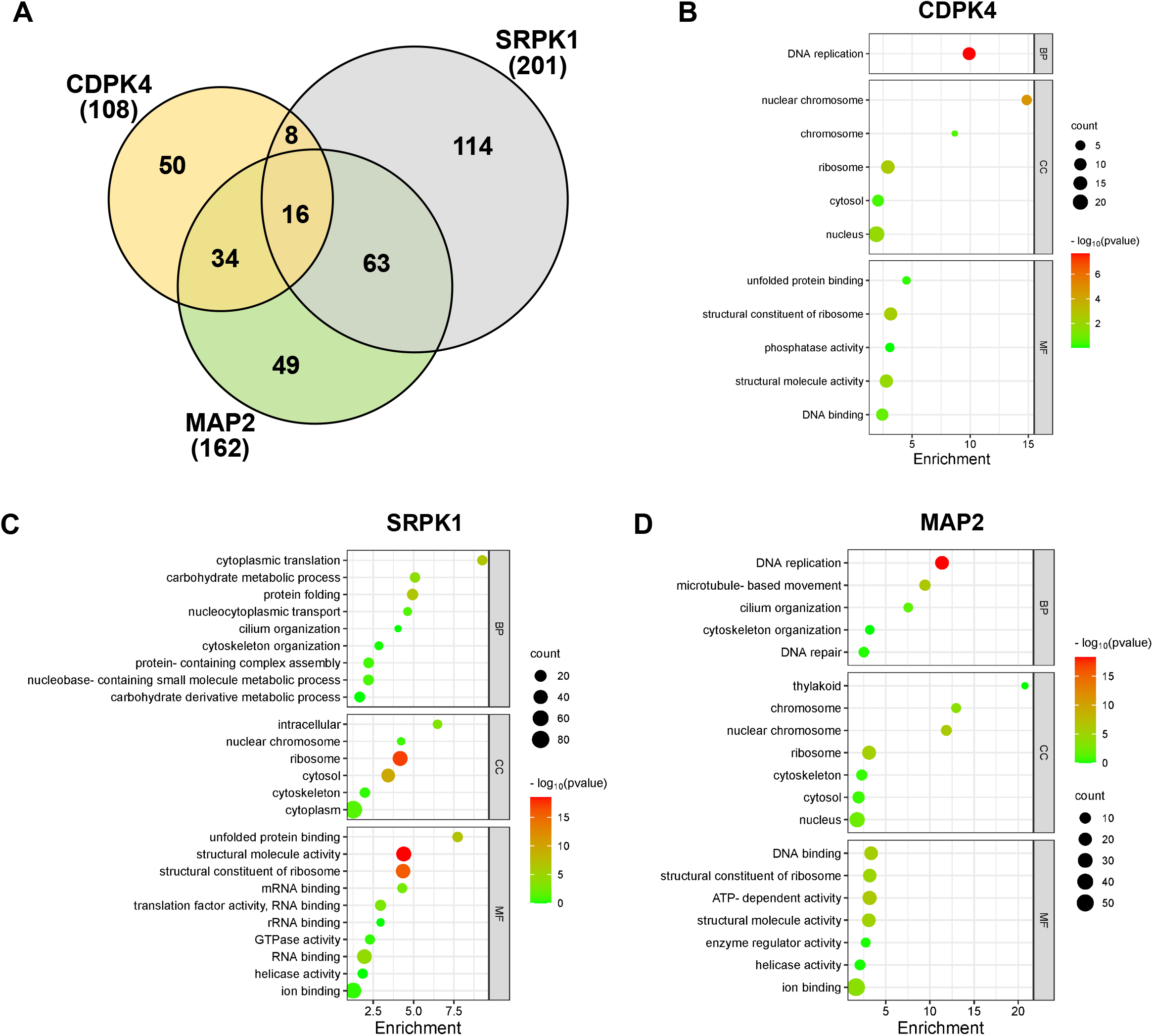
Interactome of CDPK4, SRPK1 and MAP2 during male gametogenesis. (A) Venn diagram showing common interacting partners in gametocytes activated for 6 min with some additional proteins specific to PK. (B–D) GO enrichment analysis of CDPK4-GFP (B); SRPK1-GFP (C) and MAP2-GFP (D). BP – Biological process; CC – Cellular component; MF – Molecular function. -log10 P values obtained from Bonferroni adjusted P values. See also Table 1 and Table S2-S7.

MAP2-GFP complexes were also enriched with several MTOC-associated proteins involved in axoneme development including kinesin-8B, 13 and 15 and PF16 (Figure 3D, Table 1, Table S4; Straschil et al., 2010; Zeeshan et al., 2019; Zeeshan et al., 2022). Our results are in partial agreement with those of Fang et al., 2017 and Kumar et al., 2021, who showed altered phosphopeptide abundance for all three kinesins in both PbCDPK4 and PbSRPK1 deletion mutants (Fang et al, 2017), and PfCDPK4 mutants (Kumar et al, 2021), respectively. Additionally, SMC2/4 was co-precipitated with both CDPK4 and MAP2; whereas SMC1/3 was co-precipitated with SRPK1 and MAP2. SMC1 and SMC3 are part of the cohesin complex that facilitates sister-chromatid cohesion during mitosis, whereas SMC2/4 are part of the condensin complex that regulates chromosome condensation and segregation during cell division (Uhlmann, 2016). SMC2/4 knockdown significantly affects male exflagellation and parasite transmission (Pandey et al, 2020). In yeast SMC4 activity and dynamic binding of the condensation machinery are governed by Cdk1 (Robellet et al, 2015), thus it is possible that a major regulator of condensin activity may be CDPK4 since it interacts with SMC4 (Fang et al, 2017). Our study also suggests that SMC2/4 play a role late in male gametogenesis through interaction with MAP2-GFP, possibly a role in atypical chromosome condensation through association with the MTOC (Guttery et al, 2012).

SRPK1-GFP precipitates were strongly enriched for translation pathway and 80S ribosome proteins, including eighteen 40S and twenty 60S ribosomal proteins, and several subunits of the eukaryotic translation initiation factor 3 (eIF-3) complex (Figure 3C, Table 1, Table S6). mRNA translation, translation elongation, and termination are highly coordinated processes facilitated by the 80S ribosome and an extensive array of eukaryotic initiation factors (eIFs), including eIF3 (Kramer et al, 2019). Co-precipitation of these elements was not unexpected due to SRPK1’s known role in mRNA splicing and translation (Dixit et al, 2010) and suggests that this process, including phosphorylation of the eIF3 complex (Farley et al, 2011), is highly conserved in Apicomplexa. SRPK1 also co-precipitated the cochaperone HSP40, a protein which mediates dynamic interactions of SRPK1 with the major molecular chaperones HSP70 and HSP90 in mammalian cells (Zhong et al., 2009), and proteins that were detected here (Table 1). However, unlike the findings of other studies (Dixit et al, 2010), we observed no interaction of SRPK1 with serine and arginine-rich (SR) proteins.

Finally, our data are consistent with a role of reversible protein phosphorylation as a key driver of male gametogenesis. CDPK4 and SRPK1 co-precipitated each other (Table 1), consistent with a potential feedback mechanism between these PKs, as suggested previously (Invergo et al, 2017). We also observed co-precipitation of CDPK4 and MAP2 with PPM11, a novel PP shown previously to play a role in male gametogenesis (Balestra et al, 2020). As we also found, altered phosphopeptide abundance was observed for PPM11 in CDPK4-but not SRPK1 mutants (Fang et al, 2017), which suggests that PPM11 has a role in mitotic entry and exit through dephosphorylation of CDPK4 and MAP2, respectively. CDPK4 function may also be regulated by calcineurin and PP5 (Table 1), both of which have been implicated in male gametogenesis (Philip & Waters, 2015; Zhu et al, 2019).

### Ultrastructure analysis reveals defects in MTOC formation, kinetochore dynamics, and axoneme formation in PK mutants

To determine the ultrastructural defects resulting from loss of these PKs, we used male gametocytes of CDPK4, SRPK1 and MAP2 deletion mutants (Tewari et al, 2010) activated for 8 min and examined using ultrastructure expansion microscopy (U-ExM: Gambarotto et al., 2019), and for 8 min or 15 min and examined using transmission electron microscopy (Figure 4).

**Figure 4.**
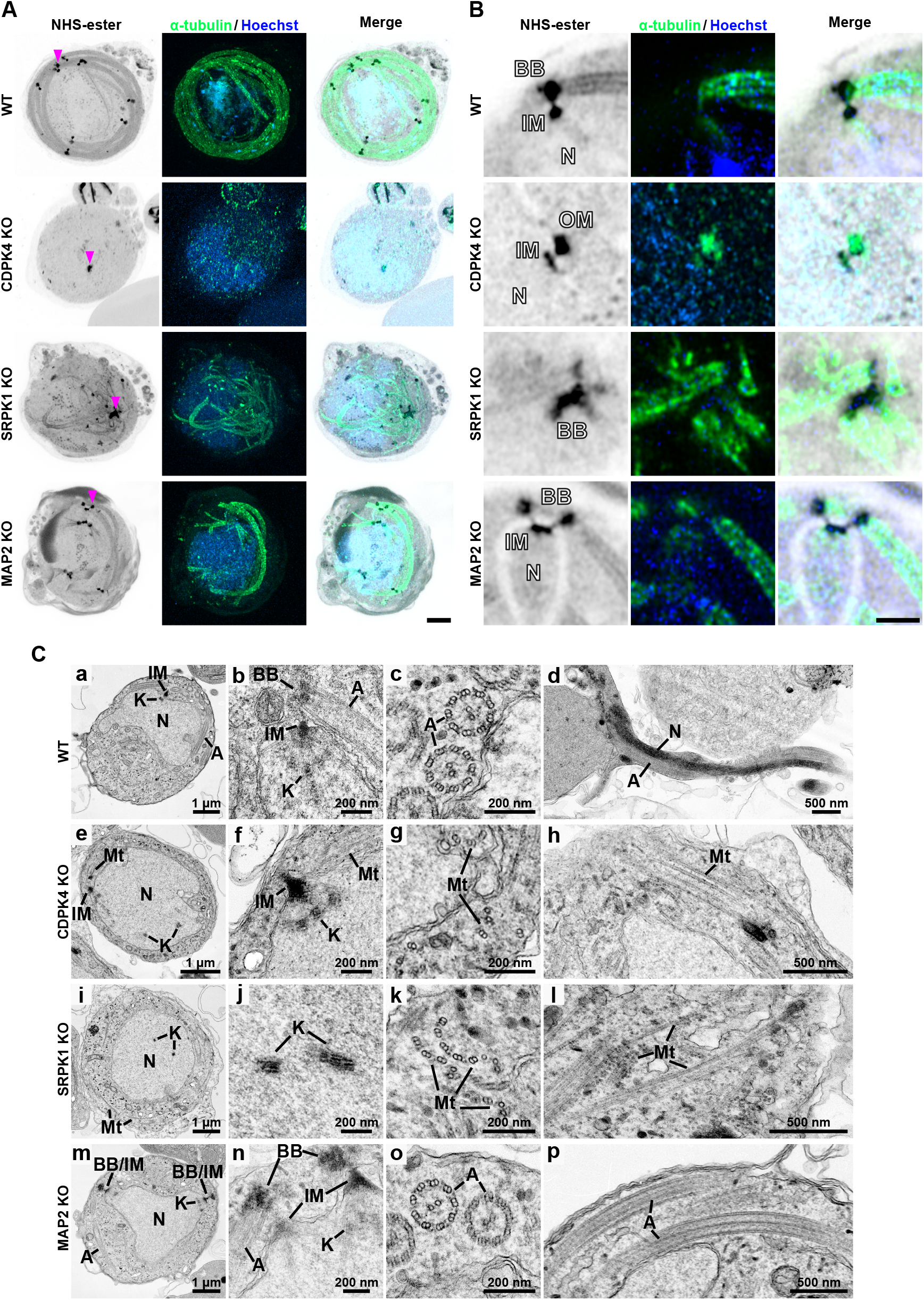
Ultrastructure analysis of wild-type and CDPK4, SRPK1 and MAP2 gene-knockout male gametocytes. (A, B) Expansion microscopy images of wild-type (WT) and CDPK4, SRPK1 and MAP2 knockout male gametocytes at 8 min post-activation. (A) Maximum intensity projections of whole-cell z-stack images labelled with NHS-ester (grey), α-tubulin (green) and Hoechst (blue). A magenta arrowhead indicates one of the MTOCs (WT, CDPK4- and MAP2 KO) or clustered basal bodies (SRPK1 KO). Scale bar = 5 µm. (B) A single z-stack image focusing on the MTOC (WT, CDPK4- and MAP2 KO) or clustered basal bodies (SRPK1 KO) region highlighted with a magenta arrowhead in (A), displaying NHS-ester (grey), α-tubulin (green), and Hoechst (blue). N: nucleus, BB: basal body, IM: inner MTOC, OM: outer MTOC. Scale bar = 2 µm. (C) Transmission electron microscopy images of WT and CDPK4-, SRPK1- and MAP2 knockout male gametocytes at 8 min (a–c, e–g, i–k, m–o) or 15 min (d, h, i, p) post-activation. N: nucleus, K: kinetochore, A: axoneme, BB: basal body, IM: inner MTOC, Mt: microtubule.

In the CDPK4 mutant, U-ExM revealed a single MTOC in the nucleus, together with a short microtubular structure extending from the MTOC (Figure 4A, B). Normal development and location of the inner MTOC, basal bodies, kinetochores axonemes and microtubules revealed in wild-type parasites by electron microscopy (Figure 4Ca-d), were disrupted in the CDPK4 mutant with some kinetochores appearing to remain within the nucleoplasm (Figure 4Ce), and several attached to the spindle radiating from the MTOC (Figure 4Cf). Dispersed doublet microtubules were present in the cytoplasm, which are likely precursors of axonemes that had failed to form (Figure 4Cf-h). These results are similar to those of Rashpa and Brochet, 2022, who had suggested that CDPK4 mutants form partial microtubules from the MTOC. This phenotype is reminiscent of that of a kinesin-13 deficient line (Zeeshan et al., 2022), consistent with our finding that CDPK4 may interacts with kinesin-13 (Table 1).

The SRPK1 mutant lacked MTOCs associated with the nucleus (Figure 4A). Kinetochores were dispersed near the centre of the nucleoplasm (Figure 4Ci, j), along with partially formed, abnormal axonemes lacking the usual 9 + 2 wild type configuration (Figure 4Cc, d, k, l). Abnormal basal bodies formed clusters away from the nucleus (Figure 4A, B). These results are again similar to the observations of Rashpa and Brochet, 2022, who showed that CDPK4 and SRPK1 are key for axoneme assembly, microtubule nucleation and development of the MTOC.

We showed previously that MAP2 deletion results in defective chromosome condensation with an arrested kinetochore during male gametogenesis (Guttery et al, 2012). In MAP2 mutant lines, the two MTOCs remained close together without separating (Figure 4A, B, Cn) unlike in wild-type cells, where they are dispersed throughout the cytoplasm (Figure 4Ca, b). Axoneme structure appeared to be similar to that in wild-type cells (Figure 4A, B), with the classical 9+2 subunit arrangement (Figure 4Cm-p). However, despite the apparent formation of intact axonemes, exflagellation was not observed. As we suggested previously (Guttery et al, 2012), MAP2 is involved in axoneme function, and impaired axoneme movement may prevent exflagellation. The MTOC separation that is also affected in the MAP2 mutant, may be another reason why gametes are unable to undergo exflagellation. These phenotypes resemble other mutant in male gametogenesis like CRK5, Kinesin-8B and kinesin-13 and CDC20 (Guttery et al, 2012; Zeeshan et al, 2019, 2022b; Balestra et al, 2020). More recently, we also observed similar chromosome segregation defects, like those seen in CDPK4 and SRPK1 knockout, during male gametogenesis in NEK1 knockdown (Zeeshan et al, 2024). This suggests that these mitotic kinases are driving the MTOC organisation and chromosome segregation. It will be interesting to study these mutants for the 3D organisation during male gametogenesis using three-dimensional electron microscopy such as serial block face scanning electron microscopy (SBF-SEM; Hair et al, 2023), in the future.

In conclusion, this study focused on three divergent protein kinases, CDPK4, SRPK1 and MAP2 that are defined as male specific because their deletion affects only the development of male gametes. We identified their location during schizogony and male gametogenesis by fluorescence live-cell imaging, and used immunoprecipitation and mass spectrometry to identify binding partners that are potential substrates. Using expansion and electron microscopy, we detailed the ultrastructural defects during male gametogenesis in cells for which each of the genes has been deleted. Our findings provide new insights into the location and function of these PKs and their substrates during male gametogenesis, which will underpin the development of drugs targeting this critical stage of the Plasmodium life cycle.

## Materials and Methods

### Ethics statement

The animal work passed an ethical review process and was approved by the United Kingdom Home Office. Work was carried out under UK Home Office Project Licenses (30/3248 and PDD2D5182) in accordance with the UK ‘Animals (Scientific Procedures) Act 1986’. Six-to eight-week-old female CD1 outbred mice from Charles River laboratories were used for all experiments.

### Generation of transgenic parasites and genotype analyses

Deletion mutants for CDPK4, SRPK1 and MAP2 were generated previously (Tewari et al, 2010). To generate the GFP-tag lines, a region of each gene downstream of the ATG start codon was amplified, ligated to p277 vector, linearised using ClaI and transfected as described previously (Guttery et al, 2012). The p277 vector contains the human dhfr cassette, conveying resistance to pyrimethamine. A schematic representation of the endogenous gene locus, the constructs and the recombined gene locus can be found in Figure S1A, with primer sequences given in Table S1. For the parasites expressing the C-terminal GFP-tagged protein, diagnostic PCR was used with primer 1 (Int primer) and primer 3 (ol492) to confirm integration of the GFP targeting construct (Figure S1B).

### Live cell imaging

To examine CDPK4-GFP, SRPK1-GFP and MAP2-GFP expression during erythrocytic stages, parasites growing in schizont culture medium were used for imaging at different stages of schizogony. Purified gametocytes were examined for GFP expression and cellular location at different time points (0, 30 s to 15 min) after activation in ookinete culture medium (Menard, 2013; Zeeshan et al, 2019). Images were captured using a 63x oil immersion objective on a Zeiss Axio Imager M2 microscope fitted with an AxioCam ICc1 digital camera.

### Generation of dual-tagged parasite lines

The GFP-tagged CDPK4, SRPK1 and MAP2 parasite lines were mixed with mCherry-tagged lines of the kinetochore marker NDC80 (Zeeshan et al, 2022b) and axoneme marker kinesin-8B (Zeeshan et al, 2019) in equal numbers and injected into mice. Mosquitoes were fed on these mice 4 to 5 days after infection when gametocytaemia was high, and oocyst development and sporozoite formation analysed at day 14 and day 21 after feeding. Infected mosquitoes were then allowed to feed on naïve mice and after 4 to 5 days the mice were examined for blood stage parasitaemia by microscopy with Giemsa-stained blood smears. Some parasites expressed both PK-GFP and NDC80-mCherry; and PK-GFP and kinesin-8B-mCherry in the resultant gametocytes, which were purified, and fluorescence microscopy images collected as described above.

### Purification of gametocytes

Parasite lines were injected into phenylhydrazine treated mice (Beetsma et al, 1998) and gametocytes enriched by sulfadiazine treatment after 2 days of infection. The blood was collected on day 4 after infection and gametocyte-infected cells were purified on a 48% v/v Nycodenz (in PBS) gradient (Nycodenz stock solution: 27.6% w/v Nycodenz in 5 mM Tris-HCl, pH 7.20, 3 mM KCl, 0.3 mM EDTA). The gametocytes were harvested from the interface and activated as described previously (Zeeshan et al, 2020).

### Immunoprecipitation and mass spectrometry

Male gametocytes of CDPK4-GFP, SRPK1-GFP and MAP2-GFP lines were used at 6 min post-activation to prepare cell lysates. WT-GFP gametocytes were used as controls. Purified parasite pellets were crosslinked using formaldehyde (10 min incubation with 1% formaldehyde, followed by 5 min incubation in 0.125 M glycine solution and three washes with phosphate-buffered saline (PBS; pH 7.5)). The cross-linked samples were solubilised in lysis buffer (10 mM Tris-HCl pH7.5, 150 mM NaCl, 0.5 mM EDTA, 0.5% Nonidet^TM^ P40 Substitute, 0.09% sodium azide with 1 x cOmplete^TM^, EDTA-free protease inhibitor cocktail (Roche)) with sonication for 1 min and lysis for 30 min on ice. Immunoprecipitation was performed using the protein lysates and a GFP-Trap_A Kit (Chromotek) following the manufacturer’s instructions. Briefly, the lysates were incubated for 2hr with GFP-Trap_A beads at 4° C with continuous rotation. Unbound proteins were washed away, and proteins bound to the GFP-Trap_A beads were digested using trypsin. The tryptic peptides were analysed by liquid chromatography–tandem mass spectrometry. Mascot (http://www.matrixscience.com/) and MaxQuant (https://www.maxquant.org/) search engines were used for mass spectrometry data analysis. Peptide and proteins having a minimum threshold of 95% were used for further proteomic analysis. The PlasmoDB database was used for protein annotation, and Gene Ontology (Ashburner et al., 2000) performed using the tool available through PlasmoDB with the following settings: computed and curated evidence allowed, use GO Slim terms, and P value cutoff = 0.05.

### Ultrastructure expansion microscopy

Sample preparation for U-ExM was performed based on previously described protocols (Rashpa & Brochet, 2022; Liffner et al, 2024). Purified and 8 min-activated gametocytes were fixed in 4% formaldehyde in PHEM buffer (60 mM PIPES, 25 mM HEPES, 10 mM EGTA, 2 mM MgCl2, pH6.9) at room temperature for 15 min. Fixed samples were attached to 10 mm round Poly-D-Lysine coated coverslips for 15 min. Coverslips were incubated overnight at 4°C in 1.4% formaldehyde (FA)/ 2% acrylamide (AA). Gelation was performed in ammonium persulfate (APS)/TEMED (10% each)/Monomer solution (23% Sodium Acrylate; 10% AA; 0,1% BIS-AA in PBS) on ice for 5 min and at 37°C for 30 min. Gels were denatured for 15 min at 37°C and for 45 min at 95°C in denaturation buffer (200 mM SDS, 200 mM NaCl, 50 mM Tris, pH9.0, in water). After denaturation, gels were incubated in distilled water overnight for complete expansion. The following day, circular gel pieces with a diameter of ∼13 mm were excised, and the gels were washed in PBS three times for 15 min to remove excess water. The gels were then incubated in blocking buffer (3% BSA in PBS) at room temperature for 30 min, incubated with monoclonal anti-α-tubulin antibody produced in mouse (T9026, Sigma-Aldrich) in blocking buffer (1:500 dilution) at 4°C overnight and washed three times for 15 min in wash buffer (0.5% v/v TWEEN-20 in PBS). The gels were incubated with 8 µg/ml Atto 594 NHS ester (Merck), 10 µg/ml Hoechst 33342 (Molecular Probes) and Alexa Fluor 488 goat anti-mouse IgG (A11001, Invitrogen) in PBS (1:500 dilution) at 37°C for 3 hours followed by three washes of 15 min each in wash buffer (blocking and all antibody incubation steps were performed with 120–160 rpm shaking). The gels were then washed three times for 15 min with wash buffer and expanded overnight in ultrapure water. The expanded gel was placed in a 35 mm glass bottom dish (MATTEK) with the 14 mm glass coated with Poly-D-Lysine and mounted with an 18 × 18 mm coverslip to prevent the gel from sliding and to avoid drifting while imaging. High-resolution confocal microscopy images were acquired using a Zeiss Celldiscoverer 7 with Airyscan using a Plan-Apochromat 50 ×/1.2NA Water objective, with 405, 488 and 561 nm lasers. Confocal z-stacks were acquired using line scanning and the following settings: 55 × 55 nm pixel size, 170 nm z-step, 2.91 µs/pixel dwell time, 850 gain and 3.5% (405 nm), 4.5% (488 nm) and 5.0% (561 nm) laser powers. The z-stack images were processed and analysed using Fiji (Version 1.54f; Schindelin et al., 2012).

### Electron microscopy

Gametocytes activated for 8 min and 15 min were fixed in 4% glutaraldehyde in 0.1 M phosphate buffer and processed for electron microscopy (Ferguson et al, 2005). Briefly, samples were post fixed in osmium tetroxide, treated en bloc with uranyl acetate, dehydrated and embedded in Spurr’s epoxy resin. Thin sections were stained with uranyl acetate and lead citrate prior to examination in a JEOL 1200EX electron microscope (JEOL Ltd, UK).

## Acknowledgements

We wish to thank Julie Rodgers for helping to maintain the insectary and other technical works and Dr Benoit Poulin for assistance. This work was supported by: MRC UK (G0900109, G0900278, MR/K011782/1) to R.T., BBSRC (BB/N017609/1) and ERC advance grant funded by UKRI Frontier Science (EP/X024776/1) to R.T., BBSRC (BB/X014681/1) to D.S.G and R.T., and BBSRC (BB/N017609/1). R.Y. is supported under grant BB/X014681/1, M.Z. is supported through grant EP/X024776/1 and BB/N017609/1. A.A.H. is supported by The Francis Crick Institute (FC001097), which receives its core funding from Cancer Research UK (FC001097), the UK Medical Research Council (FC001097), and the Wellcome Trust (FC001097). For Open Access, the authors have applied a CC BY public copyright licence to any Author Accepted Manuscript version arising from this submission.

## Author contributions

R.T. conceived and designed all experiments. R.T., D.S.G., M.Z., D.B. performed the GFP tagging experiments. R.T., and D.B. performed protein pull-down experiments. A.R.B. performed mass spectrometry. R.Y. and D.J.F.P performed ultrastructure analyses. All authors analysed the data. D.S.G., R. Y. and R.T. wrote the first draft manuscript, and all others contributed to it.

---

**Figure S1.**
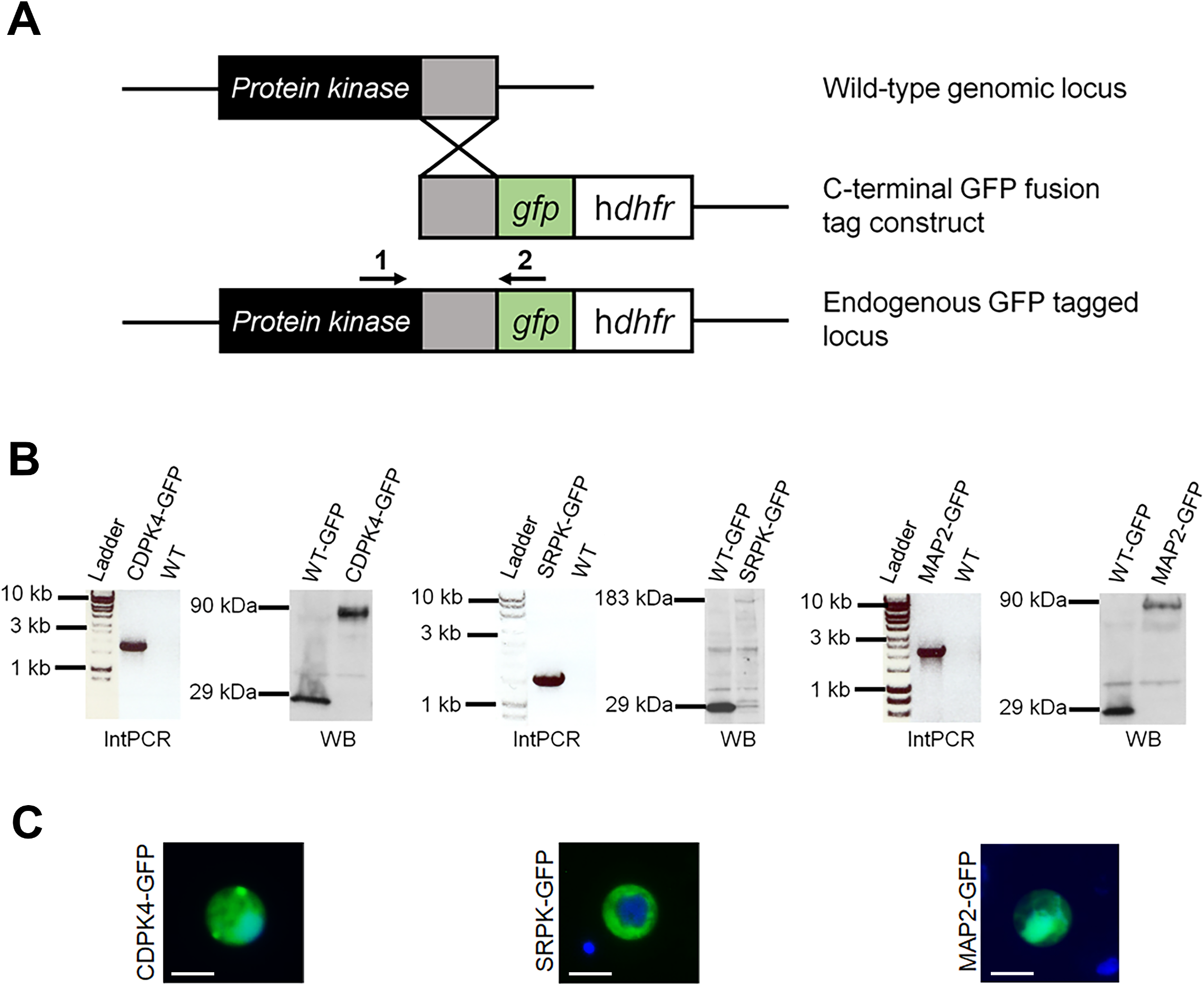
C-terminal GFP fusion tagging of the male-specific protein kinases. (A) Schematic representation for 3’-tagging of each endogenous P. berghei PK gene with gfp via single homologous recombination. Primers 1+2 used for diagnostic PCR are indicated. (B) For each PK are shown: (left) diagnostic integration PCR (IntPCR) showing band at the expected size, confirming successful integration of the tagging construct; (right) Western blot (WB) analysis using an anti-GFP antibody against control GFP (WT-GFP) and transgenic parasite protein showing bands of 29 kDa for GFP and of the expected size for the corresponding PK-GFP. (C) Images showing expression of the GFP tagged PK in male gametocytes 6 min post-activation. Scale bar = 5 µm.

**Figure S2.**
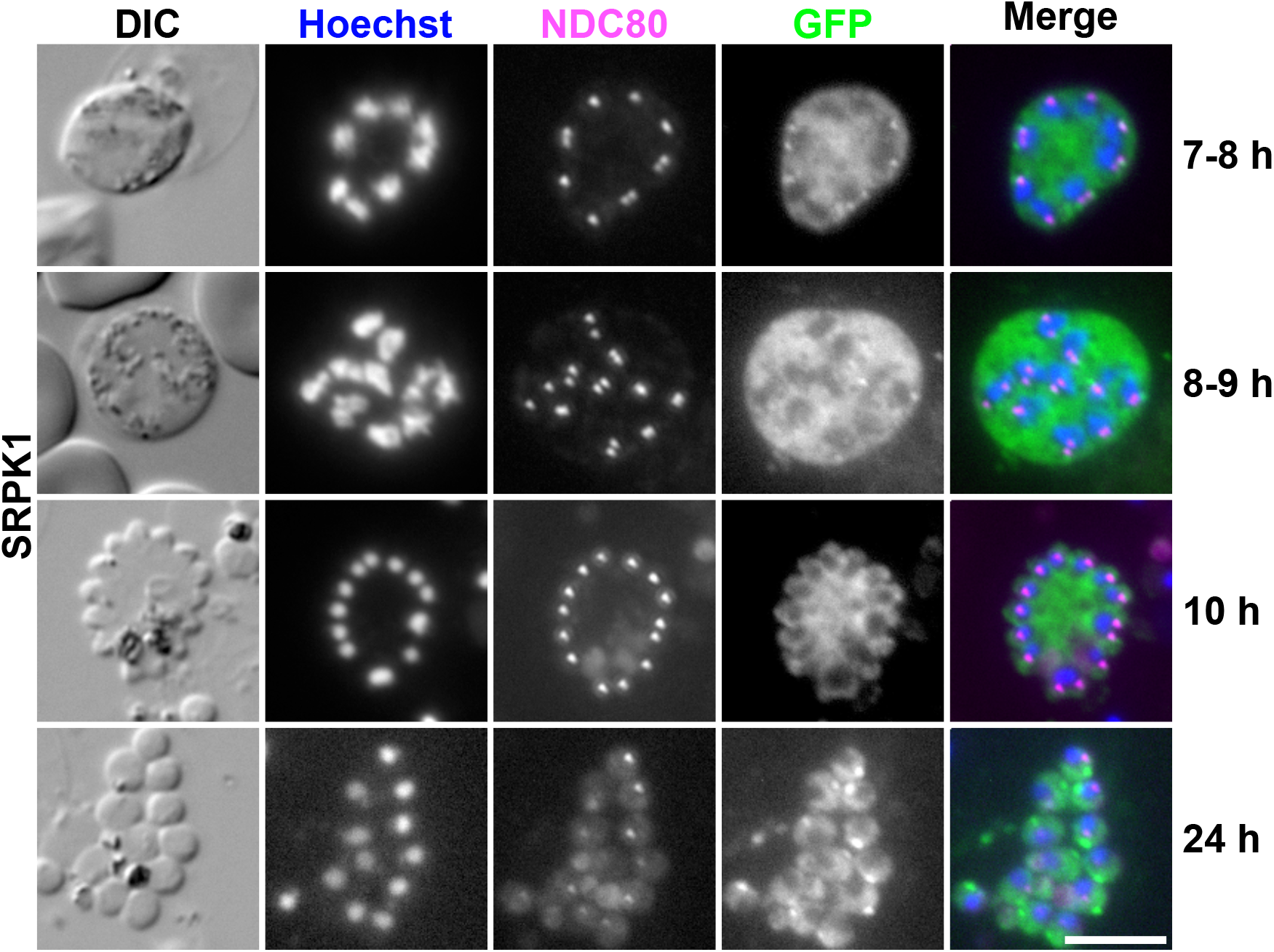
Co-localisation of SRPK1-GFP with kinetochore marker during asexual stages. Localisation of GFP-tagged SRPK1 crossed with mCh-tagged NDC80 in schizonts or merozoites. DIC: differential interference contrast. Merge shows Hoechst (blue), NDC80-mCh (magenta) and SRPK1-GFP (green). The time on the right indicates the culture duration within schizont culture medium. Scale bar = 5 µm.

## References

Balestra AC, Zeeshan M, Rea E, Pasquarello C, Brusini L, Mourier T, Subudhi AK, Klages N, Arboit P, Pandey R et al (2020) A divergent cyclin/cyclin-dependent kinase complex controls the atypical replication of a malaria parasite during gametogony and transmission. Elife 9: 1–25. doi:10.7554/eLife.56474.

Barr AR, Gergely F (2007) Aurora-A: The maker and breaker of spindle poles. J Cell Sci 120: 2987–2996. doi:10.1242/jcs.013136.

Beetsma AL, Van De Wiel TJJM, Sauerwein RW, Eling WMC (1998) Plasmodium berghei ANKA: Purification of Large Numbers of Infectious Gametocytes. Exp Parasitol 88: 69–72.

Billker O, Dechamps S, Tewari R, Wenig G, Franke-Fayard B, Brinkmann V (2004) Calcium and a calcium-dependent protein kinase regulate gamete formation and mosquito transmission in a malaria parasite. Cell 117: 503–514. doi:10.1016/S0092-8674(04)00449-0.

Carmena M, Earnshaw WC, Glover DM (2015) The dawn of Aurora kinase research: From fly genetics to the clinic. Front Cell Dev Biol 3. doi:10.3389/fcell.2015.00073.

Carmena M, Ruchaud S, Earnshaw WC (2009) Making the Auroras glow: regulation of Aurora A and B kinase function by interacting proteins. Curr Opin Cell Biol 21: 796–805. doi:10.1016/j.ceb.2009.09.008.

Chen CT, Gubbels MJ (2013) The Toxoplasma gondii centrosome is the platform for internal daughter budding as revealed by a Nek1 kinase mutant. J Cell Sci 126: 3344–3355. doi:10.1242/jcs.123364.

Chou HC, Bhalla K, Demerdesh OEL, Klingbeil O, Hanington K, Aganezov S, Andrews P, Alsudani H, Chang K, Vakoc CR et al (2021) The human origin recognition complex is essential for pre-rc assembly, mitosis, and maintenance of nuclear structure. Elife 10: 1–39. doi:10.7554/eLife.61797.

Chuang LC, Teixeira LK, Wohlschlegel JA, Henze M, Yates JR, Méndez J, Reed SI (2009) Phosphorylation of Mcm2 by Cdc7 Promotes Pre-replication Complex Assembly during Cell-Cycle Re-entry. Mol Cell 35: 206–216. doi:10.1016/j.molcel.2009.06.014.

Dixit A, Singh PK, Sharma GP, Malhotra P, Sharma P (2010) PfSRPK1, a novel splicing-related kinase from Plasmodium falciparum. Journal of Biological Chemistry 285: 38315–38323. doi:10.1074/jbc.M110.119255.

Elserafy M, Šarić M, Neuner A, Lin TC, Zhang W, Seybold C, Sivashanmugam L, Schiebel E (2014) Molecular mechanisms that restrict yeast centrosome duplication to one event per cell cycle. Current Biology 24: 1456–1466. doi:10.1016/j.cub.2014.05.032.

Fang H, Klages N, Baechler B, Hillner E, Yu L, Pardo M, Choudhary J, Brochet M (2017) Multiple short windows of calcium-dependent protein kinase 4 activity coordinate distinct cell cycle events during Plasmodium gametogenesis. Elife 6: 1–23. doi:10.7554/eLife.26524.001.

Farley AR, Powell DW, Weaver CM, Jennings JL, Link AJ (2011) Assessing the components of the eIF3 complex and their phosphorylation status. J Proteome Res 10: 1481–1494. doi:10.1021/pr100877m.

Ferguson DJP, Henriquez FL, Kirisits MJ, Muench SP, Prigge ST, Rice DW, Roberts CW, McLeod RL (2005) Maternal inheritance and stage-specific variation of the apicoplast in Toxoplasma gondii during development in the intermediate and definitive host. Eukaryot Cell 4: 814–826. doi:10.1128/EC.4.4.814-826.2005.

Fry AM, Bayliss R, Roig J (2017) Mitotic regulation by NEK kinase networks. Front Cell Dev Biol 5. doi:10.3389/fcell.2017.00102.

Fulcher LJ, Sapkota GP (2020) Mitotic kinase anchoring proteins: the navigators of cell division. Cell Cycle 19: 505–524. doi:10.1080/15384101.2020.1728014.

Gaji RY, Sharp AK, Brown AM (2021) Protein kinases in Toxoplasma gondii. Int J Parasitol 51: 415–429. doi:10.1016/j.ijpara.2020.11.006.

Gambarotto D, Zwettler FU, Le Guennec M, Schmidt-Cernohorska M, Fortun D, Borgers S, Heine J, Schloetel JG, Reuss M, Unser M et al (2019) Imaging cellular ultrastructures using expansion microscopy (U-ExM). Nat Methods 16: 71–74. doi:10.1038/s41592-018-0238-1.

Guttery DS, Ferguson DJP, Poulin B, Xu Z, Straschil U, Klop O, Solyakov L, Sandrini SM, Brady D, Nieduszynski CA et al (2012) A putative homologue of CDC20/CDH1 in the malaria parasite is essential for male gamete development. PLoS Pathog 8. doi:10.1371/journal.ppat.1002554.

Guttery DS, Zeeshan M, Ferguson DJP, Holder AA, Tewari R (2022) Division and Transmission: Malaria Parasite Development in the Mosquito. Annu Rev Microbiol 76: 113–134. doi:10.1146/annurev-micro-041320-010046.

Guttery DS, Zeeshan M, Holder AA, Tewari R (2024) The molecular mechanisms driving Plasmodium cell division. Biochem Soc Trans 52: 593–602. doi:10.1042/BST20230403.

Hadders MA, Lens SMA (2022) Changing places: Chromosomal Passenger Complex relocation in early anaphase. Trends Cell Biol 32: 165–176. doi:10.1016/j.tcb.2021.09.008.

Hair M, Moreira-Leite F, Ferguson DJP, Zeeshan M, Tewari R, Vaughan S (2023) Atypical flagella assembly and haploid genome coiling during male gamete formation in Plasmodium. Nat Commun 14. doi:10.1038/s41467-023-43877-w.

Honda R, Körner R, Nigg EA (2003) Exploring the functional interactions between Aurora B, INCENP, and survivin in mitosis. Mol Biol Cell 14: 3325–3341. doi:10.1091/mbc.E02-11-0769.

Invergo BM, Brochet M, Yu L, Choudhary J, Beltrao P, Billker O (2017) Sub-minute Phosphoregulation of Cell Cycle Systems during Plasmodium Gamete Formation. Cell Rep 21: 2017–2029. doi:10.1016/j.celrep.2017.10.071.

Ito D, Bettencourt-Dias M (2018) Centrosome remodelling in evolution. Cells 7. doi:10.3390/cells7070071.

Joubes J, Chevalier C, Dudits D, Heberle-Bors E, Inze D, Umeda M, Renaudin J-P (2000) CDK-related protein kinases in plants. Plant Mol Biol 43: 607–620.

Kramer G, Shiber A, Bukau B (2019) Annual Review of Biochemistry Mechanisms of Cotranslational Maturation of Newly Synthesized Proteins. Annu Rev Biochem 88: 337–364. doi:10.1146/annurev-biochem-013118.

Kucharski TJ, Hards R, Vandal SE, Abad MA, Jeyaprakash AA, Kaye E, Al-Rawi A, Ly T, Godek KM, Gerber SA et al (2022) Small changes in phospho-occupancy at the kinetochore–microtubule interface drive mitotic fidelity. Journal of Cell Biology 221. doi:10.1083/jcb.202107107.

Kumar S, Gargaro OR, Kappe SHI (2022) Plasmodium falciparum CRK5 Is Critical for Male Gametogenesis and Infection of the Mosquito. mBio 13. doi:10.1128/mbio.02227-22.

Kumar S, Haile MT, Hoopmann MR, Tran LT, Michaels SA, Morrone SR, Ojo KK, Reynolds LM, Kusebauch U, Vaughan AM et al (2021) Plasmodium falciparum Calcium-Dependent Protein Kinase 4 is Critical for Male Gametogenesis and Transmission to the Mosquito Vector. mBio 12. doi:10.1128/mBio.02575-21.

Lee KY, Bang SW, Yoon SW, Lee SH, Yoon JB, Hwang DS (2012) Phosphorylation of ORC2 protein dissociates origin recognition complex from chromatin and replication origins. Journal of Biological Chemistry 287: 11891–11898. doi:10.1074/jbc.M111.338467.

Liffner B, Alves E Silva TL, Vega-Rodriguez J, Absalon S (2024) Mosquito Tissue Ultrastructure-Expansion Microscopy (MoTissU-ExM) enables ultrastructural and anatomical analysis of malaria parasites and their mosquito. bioRxiv. doi:10.1101/2024.04.17.589980.

Menard R (2013) Malaria: Methods and Protocols (Humana Press).

Nilsson J (2019) Protein phosphatases in the regulation of mitosis. Journal of Cell Biology 218: 395–409. doi:10.1083/jcb.201809138.

Ong JY, Bradley MC, Torres JZ (2020) Phospho-regulation of mitotic spindle assembly. Cytoskeleton 77: 558–578. doi:10.1002/cm.21649.

Panchal NK, Evan Prince S (2023) The NEK family of serine/threonine kinases as a biomarker for cancer. Clin Exp Med 23: 17–30. doi:10.1007/s10238-021-00782-0.

Pandey R, Abel S, Boucher M, Wall RJ, Zeeshan M, Rea E, Freville A, Lu XM, Brady D, Daniel E et al (2020) Plasmodium Condensin Core Subunits SMC2/SMC4 Mediate Atypical Mitosis and Are Essential for Parasite Proliferation and Transmission. Cell Rep 30: 1883-1897.e6. doi:10.1016/j.celrep.2020.01.033.

Philip N, Waters AP (2015) Conditional degradation of plasmodium calcineurin reveals functions in parasite colonization of both host and vector. Cell Host Microbe 18: 122–131. doi:10.1016/j.chom.2015.05.018.

Rashpa R, Brochet M (2022) Expansion microscopy of Plasmodium gametocytes reveals the molecular architecture of a bipartite microtubule organisation centre coordinating mitosis with axoneme assembly. PLoS Pathog 18. doi:10.1371/journal.ppat.1010223.

Robellet X, Thattikota Y, Wang F, Wee TL, Pascariu M, Shankar S, Bonneil Eé, Brown CM, D’amours D (2015) A high-sensitivity phospho-switch triggered by Cdk1 governs chromosome morphogenesis during cell division. Genes Dev 29: 426–439. doi:10.1101/gad.253294.114.

Schindelin J, Arganda-Carreras I, Frise E, Kaynig V, Longair M, Pietzsch T, Preibisch S, Rueden C, Saalfeld S, Schmid B et al (2012) Fiji: An open-source platform for biological-image analysis. Nat Methods 9: 676–682. doi:10.1038/nmeth.2019.

Seybold C, Shiebel E (2013) Spindle pole bodies. Current Biology 23. doi:10.1016/j.cub.2013.07.024.

Sheu YJ, Stillman B (2006) Cdc7-Dbf4 Phosphorylates MCM Proteins via a Docking Site-Mediated Mechanism to Promote S Phase Progression. Mol Cell 24: 101–113. doi:10.1016/j.molcel.2006.07.033.

Sinden RE (2015) The cell biology of malaria infection of mosquito: Advances and opportunities. Cell Microbiol 17: 451–466. doi:10.1111/cmi.12413.

Straschil U, Talman AM, Ferguson DJP, Bunting KA, Xu Z, Bailes E, Sinden RE, Holder AA, Smith EF, Coates JC et al (2010) The armadillo repeat protein PF16 is essential for flagellar structure and function in Plasmodium male gametes. PLoS One 5. doi:10.1371/journal.pone.0012901.

Strumillo MJ, Oplová M, Viéitez C, Ochoa D, Shahraz M, Busby BP, Sopko R, Studer RA, Perrimon N, Panse VG et al (2019) Conserved phosphorylation hotspots in eukaryotic protein domain families. Nat Commun 10. doi:10.1038/s41467-019-09952-x.

Suvorova ES, Francia M, Striepen B, White MW (2015) A Novel Bipartite Centrosome Coordinates the Apicomplexan Cell Cycle. PLoS Biol 13. doi:10.1371/journal.pbio.1002093.

Tewari R, Dorin D, Moon R, Doerig C, Billker O (2005) An atypical mitogen-activated protein kinase controls cytokinesis and flagellar motility during male gamete formation in a malaria parasite. Mol Microbiol 58: 1253–1263. doi:10.1111/j.1365-2958.2005.04793.x.

Tewari R, Straschil U, Bateman A, Böhme U, Cherevach I, Gong P, Pain A, Billker O (2010) The systematic functional analysis of plasmodium protein kinases identifies essential regulators of mosquito transmission. Cell Host Microbe 8: 377–387. doi:10.1016/j.chom.2010.09.006.

Uhlmann F (2016) SMC complexes: From DNA to chromosomes. Nat Rev Mol Cell Biol 17: 399– 412. doi:10.1038/nrm.2016.30.

Vagnarelli P (2021) Back to the new beginning: Mitotic exit in space and time. Semin Cell Dev Biol 117: 140–148. doi:10.1016/j.semcdb.2021.03.010.

Vaughan S, Gull K (2016) Basal body structure and cell cycle - dependent biogenesis in Trypanosoma brucei. Cilia: 1–7. doi:10.1186/s13630-016-0023-7.

Vertii A, Hehnly H, Doxsey S (2016) The centrosome, a multitalented renaissance organelle. Cold Spring Harb Perspect Biol 8. doi:10.1101/cshperspect.a025049.

Wu J, Akhmanova A (2017) Microtubule-Organizing Centers. Annu Rev Cell Dev Biol 33: 51–75. doi:10.1146/annurev-cellbio-100616.

Zeeshan M, Brady D, Markus R, Vaughan S, Ferguson D, Holder AA, Tewari R (2022a) Plasmodium SAS4: basal body component of male cell which is dispensable for parasite transmission. Life Sci Alliance 5. doi:10.26508/lsa.202101329.

Zeeshan M, Ferguson DJP, Abel S, Burrrell A, Rea E, Brady D, Daniel E, Delves M, Vaughan S, Holder AA et al (2019) Kinesin-8B controls basal body function and flagellum formation and is key to malaria transmission. Life Sci Alliance 2. doi:10.26508/lsa.201900488.

Zeeshan M, Pandey R, Ferguson DJP, Tromer EC, Markus R, Abel S, Brady D, Daniel E, Limenitakis R, Bottrill AR et al (2020) Real-time dynamics of Plasmodium NDC80 reveals unusual modes of chromosome segregation during parasite proliferation. J Cell Sci 134. doi:10.1242/JCS.245753.

Zeeshan M, Rashpa R, Ferguson DJ, Mckeown G, Nugmanova R, Subudhi AK, Beyeler R, Pashley SL, Markus R, Brady D et al (2024) Plasmodium NEK1 coordinates MTOC organisation and kinetochore attachment during rapid mitosis in male gamete formation. PLoS Biol 22: e3002802. doi:10.1371/journal.pbio.3002802.

Zeeshan M, Rashpa R, Ferguson DJP, Abel S, Chahine Z, Brady D, Vaughan S, Moores CA, Le Roch KG, Brochet M et al (2022b) Genome-wide functional analysis reveals key roles for kinesins in the mammalian and mosquito stages of the malaria parasite life cycle. PLoS Biol 20. doi:10.1371/journal.pbio.3001704.

Zeeshan M, Rea E, Abel S, Vukušić K, Markus R, Brady D, Eze A, Rashpa R, Balestra AC, Bottrill AR et al (2023) Plasmodium ARK2 and EB1 drive unconventional spindle dynamics, during chromosome segregation in sexual transmission stages. Nat Commun 14. doi:10.1038/s41467-023-41395-3.

Zhong XY, Ding JH, Adams JA, Ghosh G, Fu XD (2009) Regulation of SR protein phosphorylation and alternative splicing by modulating kinetic interactions of SRPK1 with molecular chaperones. Genes Dev 23: 482–495. doi:10.1101/gad.1752109.

Zhu X, Sun L, He Y, Wei H, Hong M, Liu F, Liu Q, Cao Y, Cui L (2019) Plasmodium berghei serine/threonine protein phosphatase PP5 plays a critical role in male gamete fertility. Int J Parasitol 49: 685–695. doi:10.1016/j.ijpara.2019.03.007.

